# The PAF1 complex cell-autonomously promotes oogenesis in *Caenorhabditis elegans*

**DOI:** 10.1101/2021.08.18.456829

**Authors:** Yukihiro Kubota, Natsumi Ota, Hisashi Takatsuka, Takuma Unno, Shuichi Onami, Asako Sugimoto, Masahiro Ito

**Affiliations:** Department of Bioinformatics, College of Life Sciences, Ritsumeikan University, 1-1-1 Nojihigashi, Kusatsu, Shiga, Japan; Advanced Life Sciences Program, Graduate School of Life Sciences, Ritsumeikan University, 1-1-1 Nojihigashi, Kusatsu, Shiga, Japan; RIKEN Center for Biosystems Dynamics Research, 2-2-3, Minatojima-minamimachi, Chuo-ku, Kobe, Japan; Laboratory of Developmental Dinamics, Graduate School of Life Sciences, Tohoku University, 2-1-1 Katahira, Sendai, Miyagi, Japan

## Abstract

The RNA polymerase II-associated factor 1 complex (PAF1C) is a protein complex that consists of LEO1, RTF1, PAF1, CDC73, and CTR9, and has been shown to be involved in Pol II-mediated transcriptional and chromatin regulation. Although it has been shown to regulate a variety of biological processes, the precise role of the PAF1C during germ line development has not been clarified. In this study, we found that reduction in the function of the PAF1C components, LEO-1, RTFO-1, PAFO-1, CDC-73, and CTR-9, in *Caenorhabditis elegans* affects cell volume expansion of oocytes. Defects in oogenesis were also confirmed using an oocyte maturation marker, OMA-1::GFP. While four to five OMA-1::GFP-positive oocytes were observed in wild-type animals, their numbers were significantly decreased in *pafo-1* mutant and *leo-1(RNAi), cdc-73(RNAi)*, and *pafo-1(RNAi)* animals. Expression of a functional PAFO-1::mCherry transgene in the germline significantly rescued the oogenesis-defective phenotype of the *pafo-1* mutants, suggesting that expression of the PAF1C in germ cells is required for oogenesis. Notably, overexpression of OMA-1::GFP partially rescued the oogenesis defect in the *pafo-1* mutants. Based on our findings, we propose that the PAF1C promotes oogenesis in a cell-autonomous manner by positively regulating the expression of genes involved in oocyte maturation.

## Introduction

During animal development, spatiotemporal regulation of gene transcription is essential for precise regulation of cell behavior. To precisely regulate gene transcription, the recruitment and activation of RNA polymerase II (Pol II) to the transcriptional target is required. In addition, chromatin remodeling affects DNA accessibility during transcription through epigenetic modification of nucleosomes. The polymerase associated factor 1(PAF1) complex, or PAF1C, is a highly conserved protein complex in eukaryotes, which is involved in multiple aspects of Pol II-mediated transcriptional regulation, including transcriptional elongation, 3′-end processing, and epigenetic modification. Moreover, the PAF1C is involved in the post-transcriptional step of gene expression and translational regulation via its interaction with the regulatory sequences of mRNAs [1, 2].

The PAF1C was originally identified in *Saccharomyces cerevisiae* as an RNA pol II interactor [3-5]. It consists of five subunits (Leo1, Rtf1, Paf1/pancreatic differentiation, Cdc73/parafibromin, and Ctr9) [6, 7]. Although PAF1C is not essential for the viability of *S. cerevisiae*, depletion or mutation of the PAF1 subunits causes severe developmental disorders during the development of somite, neural crest, neuron, heart, and craniofacial cartilage in zebrafish [8-11]. Additionally, the PAF1C affects Notch, Wnt, and Hedgehog signaling [8, 12-14]. The PAF1C has also been reported to regulate the proliferation, differentiation, morphology, cell migration, epidermal morphogenesis, mitophagy, maintenance of stem cells, and tumorigenesis [4, 10, 15-23]. However, the functional importance of the PAF1C in germ cell development has not yet been explored.

The development of the nematode, *Caenorhabditis elegans*, is highly reproducible, which makes it a reliable model organism for analyzing the regulatory mechanism of development. The hermaphrodite gonad of this nematode temporally produces sperm at the late larval stage, which are stored in the spermatheca, and subsequently produces oocytes during the adult stage. During this process, spatiotemporal regulation of gene expression, cell proliferation, cell differentiation, cell shape change, cell growth, and meiotic progression occurs [24-26]. However, the mechanism of oogenesis has not been fully elucidated.

In this study, we found that all the PAF1C components are involved in promoting the expansion of cell volume of oocytes, and that the expression of OMA-1, a CCCH-type zinc finger protein involved in oocyte maturation, is promoted by the PAF1C in a cell-autonomous manner.

## Materials and Methods

### *C. elegans* strains

*C. elegans* strains used in this study were derived from the wild-type (WT) Bristol strain N2 [27]. Worms were incubated at 20 °C, except those that were fed RNAi bacteria and were maintained at 22 °C.

The *leo-1* locus encodes a predicted polypeptide of 430 amino acids (aa), and the *gk1081* allele (isolated by the *C. elegans* Gene Knockout Consortium) deleted 627 bp that would result in a C-terminally truncated protein of 137 aa (intrinsic 132 aa with an extra 5 aa) [20]. The *rtfo-1* locus encodes a predicted polypeptide of 613 aa, and the *tm5670* allele (isolated by the National Bioresource Project Japan) deleted 361 bp that would result in a C-terminally truncated protein product of 349 aa (intrinsic 314 aa with an extra 35 aa). The *pafo-1* locus encodes a predicted polypeptide of 425 aa, and the *tm13447* allele (isolated by the National Bioresource Project, Japan) deleted 83 bp that would result in a C-terminally truncated protein product of 288 aa (intrinsic 285 aa with an extra 3 aa). The *leo-1(gk1081)* mutant has been shown to produce reduced amounts of C-terminally truncated proteins [20]. The C-terminus of RTF1 in yeast has been shown to be required for its efficient anchoring to the PAF1C; *rtfo-1(tm5670)* is expected to be a complex formation-defective mutant [28]. We also used the following alleles for construction of mutants: *tjIs57[pie-1p::mCherry::H2B::pie-1 3′-UTR + unc-119(+)]* [29], *bkcSi11[oma-1p::oma-1::GFP::oma-1 3’-UTR, NeoR] IV, bkcSi12[pie-1p::pafo-1::mCherry::pie-1 3’-UTR, NeoR], bkcSi13[pie-1p::pafo-1::mCherry::pie-1 3’-UTR, NeoR]* (this work), *tjIs280[pafo-1p::pafo-1::mCherry::pafo-1 3′-UTR, Cbr-unc119(+)], tjIs308[leo-1p::GFP::leo-1::leo-1 3′-UTR, Cbr-unc-119(+)]* [20], *TmC3V[TmIs1230]*, and *TmC5 IV[tmIs1220]* [30].

To obtain *leo-1(gk1081)* homozygote hermaphrodites, *leo-1(gk1081)* was balanced with *TmC5 IV[tmIs1220]*, and Venus-negative homozygote progeny was scored. To obtain *rtfo-1(tm5670) and pafo-1(tm13347)* homozygote hermaphrodites, *rtfo-1(tm5670)* and *pafo-1(tm13347)* were balanced with *TmC3V[TmIs1230]*, and mCherry-negative homozygote progeny was scored.

The strains used in this work are listed in S1 Table.

### Plasmid construction

The plasmids used in this study are listed in S2 Table. A miniMos backbone vector, denoted as pYK13, was constructed by inserting a 376 bp fragment containing a multicloning site into a StuI site in pCFJ910. To construct the targeting vectors, the following fragments were amplified and individually subcloned into pYK13 at the NotI and AscI sites. To construct transgenes that expressed GFP-fusion proteins from putative endogenous 5′-cis regulatory regions of *oma-1*, the genomic fragments, *oma-1::GFP* derived from the *oma-1* regulatory region (2860 bp), the coding region, and the *oma-1* 3′-UTR (2935 bp), were PCR amplified and then fused with GFP.

For germ cell-specific expression experiments, the *pafo-1* genome was subcloned into a carboxyl-terminal mCherry-fusion protein expression vector, which has cis regulatory regions of *pie-1*, pYK229, a modified vector derived from pYK13.

### Strain construction for rescue experiments

Transgenic worms were prepared by microinjection of the target gene [31]. Strains that expressed *oma-1::GFP* under putative endogenous 5′-cis regulatory regions and 3′-cis regulatory region of *oma-1*, and strains that expressed PAFO-1::mCherry under the germ cell-specific regulatory regions of *pie-1* were used for miniMos methods (see below). Single-copy transgenic-insertion worms were generated using the miniMos method [32] for genomic GFP/mCherry-fusion expression and tissue-specific rescue experiments with the wild-type as the host strain. For microinjections, the following mixtures were used: 10 μg/mL each of GFP/mCherry-tagged miniMos-target transgene (*oma-1p::oma-1::GFP:: oma-1 3’-UTR + NeoR* plasmid pYK29, *pie-1-1p::pafo-1::mCherry:: pie-1 3’-UTR + NeoR* plasmid pYK232); transposase pCFJ601, 50 μg/mL; injection markers *Prab-3::mCherry::unc-54 3’-UTR* plasmid pGH8, 10 μg/mL; *Pmyo-2::mCherry::unc-54 3’-UTR* plasmid pCFJ90, 2.5 μg/mL; *Pmyo-3::mCherry::unc-54 3’-UTR* plasmid pCFJ104, 5 μg/mL; and pBluescript II KS(–), 30 μg/mL; negative selection marker *Phsp-16*.*41::peel-1::tbb-2 3’-UTR* plasmid pMA122, 10 μg/mL.

The integrated alleles, *tjIs280[pafo-1p::pafo-1::mCherry::pafo-1 3’-UTR, Cbr-unc119(+)], bkcSi12[pie-1p::pafo-1::mCherry::pie-1 3’-UTR, NeoR]*, and *bkcSi13[pie-1p::pafo-1::mCherry::pie-1 3’-UTR, NeoR]* were introduced to the *pafo-1(tm13347)* mutant or *bkcSi11 [oma-1p::oma-1::GFP::oma-1 3’-UTR, NeoR]*; *pafo-1(tm13347)* mutant background. Day 1 adult worms were used to score the oogenesis defects.

### Feeding RNAi

The worms were fed on RNAi-feeding plates as previously described [33]. Full-length *leo-1, rtfo-1, pafo-1*, and *cdc-73* cDNAs and 1000 bp *ctr-9* (1st–1000th coding region) cDNA were isolated from a *C. elegans* cDNA library and inserted into the feeding RNAi vector, L4440. An L4440 vector lacking an insert was used as a *control(RNAi)*. After confirming that each inserted sequence was correct, the feeding vectors were individually transformed into *Escherichia coli* HT115 (DE3) samples, which were then seeded separately onto plates of nematode growth medium agar containing Luria-Bertani medium and 50 μg/mL ampicillin, and incubated overnight at 37 °C. Thereafter, each culture was seeded onto a 60 mm feeding agar plate containing 50 μg/mL ampicillin and 1 mM isopropyl *β*-D-1-thiogalactopyranoside and incubated at 23 °C for 2 days. L4-stage worms were transferred to a feeding plate and cultured at 22 °C. Phenotypes of F1 worms were determined at the day 1 adult stage.

### Microscopy

Fluorescence and differential interference contrast; DIC microscopy procedures were performed using an Olympus BX63 microscope with an ORCA-Spark camera (Hamamatsu Photonics) and UPlanSApo X60 water NA 1.20 or UPlanXapo X40 NA 0.95 objective lens. The microscope system was controlled using the cellSens Dimension software (Olympus). Images were processed using the ImageJ (NIH) or Adobe Photoshop 2021 software.

### Statistical analyses

The *P-*value for the Fisher’s exact test for the percentage of animals with oogenesis defects in *leo-1(ok1018), rtfo-1(tm5670)*, and *pafo-1(tm13347)* were calculated for comparison with wild-type animals. The *P-*value for the Fisher’s exact test for the percentage of animals with oogenesis defects in *leo-1(RNAi), rtfo-1(RNAi), pafo-1(RNAi), cdc-73(RNAi)*, and *ctr-9(RNAi)* were calculated for comparison with *control(RNAi)* animals. For the OMA-1::GFP overexpression experiment, the *P-*value for the Fisher’s exact test for the percentage of animals with oogenesis defects in *pafo-1(tm13347);bkcSi11[oma-1p::oma-1::GFP::oma-1 3’-UTR, NeoR]* was calculated for comparison with *pafo-1(tm13347)* animals.

The *P-*value for the Student’s *t*-test of the relative expression level of GFP::LEO-1 at the distal gonad-arm region in *leo-1(RNAi);tjIs308[leo-1p::GFP::leo-1::leo-1 3’-UTR, Cbr-unc-119(+)]* was calculated for comparison with the *control(RNAi);tjIs308[leo-1p::GFP::leo-1::leo-1 3’-UTR, Cbr-unc-119(+)]* animals. The *P-*value for the Student’s *t*-test of the relative expression level of PAFO-1::mCherry at the distal gonad-arm region in *pafo-1(RNAi);tjIs280[pafo-1p::pafo-1::mCherry::pafo-1 3’-UTR, Cbr-unc119(+)]* was calculated for comparison with the *control(RNAi); tjIs280[pafo-1p::pafo-1::mCherry::pafo-1 3’-UTR, Cbr-unc119(+)]* animals. For rescue experiments, the *P-*value for the Student’s *t*-test of the number of OMA-1::GFP positive oocytes in *pafo-1(tm13347);tjIs280[pafo-1p::pafo-1::mCherry::pafo-1 3’-UTR];bkcSi11[oma-1p::oma-1::GFP::oma-1 3’-UTR, NeoR], pafo-1(tm13347);bkcSi12[pie-1p::pafo-1::mCherry::pie-1 3’-UTR];bkcSi11[oma-1p::oma-1::GFP::oma-1 3’-UTR, NeoR], pafo-1(tm13347);bkcSi13[pie-1p::pafo-1::mCherry::pie-1 3’-UTR];bkcSi11[oma-1p::oma-1::GFP::oma-1 3’-UTR, NeoR]* were calculated for comparison with the *pafo-1(tm13347);bkcSi11[oma-1p::oma-1::GFP::oma-1 3’-UTR, NeoR]* animals.

### Data availability

All data and samples described in this work will be freely provided upon request.

## Results

### The PAF1C is essential for oogenesis

To analyze whether the PAF1C is involved in germ cell development in *C. elegans*, we observed germ cell development of the posterior gonads in day 1 adults of the PAF1C mutants by DIC microscopy (Fig 1). Compared with the wild-type animals, the cell volume expansion of oocytes was insufficient in the PAF1C mutants, *leo-1(gk1081), rtfo-1(tm5670)*, and *pafo-1(tm13447)* mutants, although the penetrance of *leo-1(gk1081)* was lower than that of the other mutants (Fig 1A–1E and 1P). An integrated transgene, *pafo-1::mCherry*, expressed by the *pafo-1* regulatory element rescued the oogenesis defect of the *pafo-1(tm13447)* deletion mutant (Fig 1F and 1P). Similar to the observations of deletion mutants, although the penetrance of cell volume expansion defect of the *leo-1(RNAi)* was relatively low, all the five RNAi-knockdown animals of the PAF1C components exhibited cell volume expansion defects (Fig 1G–1M and 1P). The efficiency of both *leo-1(RNAi)* and *pafo-1(RNAi)* was over 90% (Fig. 1N and 1O; S1 Fig; S2 Fig), as measured by the fluorescent signal of GFP::LEO-1 and PAFO-1::mCherry, respectively, at the distal region of the gonad. These results suggest that the PAF1C is essential for oogenesis, and that the contribution of LEO-1 to the PAF1C function may be the lowest among the PAF1C components.

**Fig 1.**
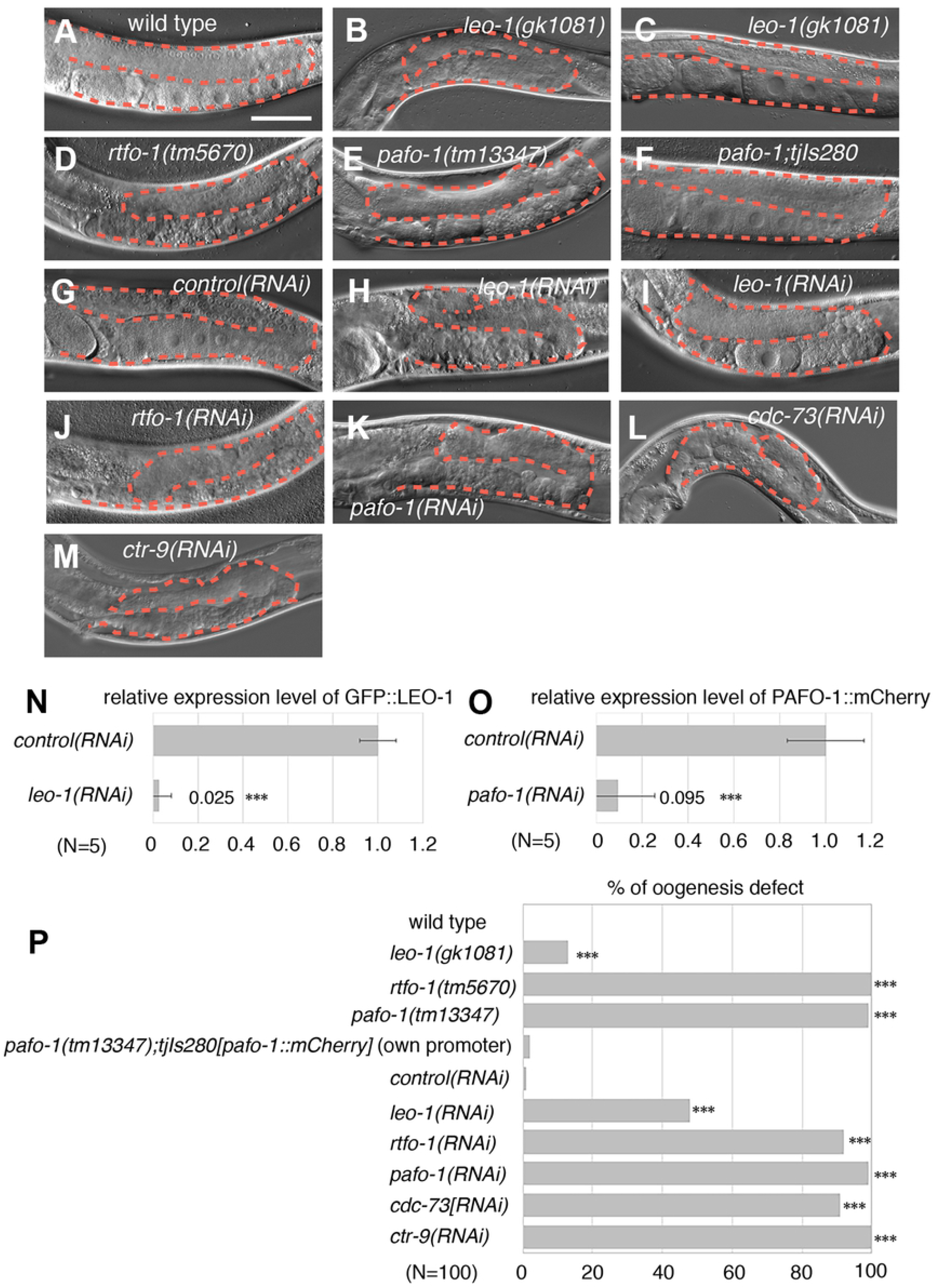
The PAF1C is Essential for Oogenesis. (A–M) Germ cell development of wild-type (A), *leo-1(gk1018)* (B, C), *rtfo-1(tm5670)* (D), *pafo-1(tm13347)* (E), *pafo-1(tm13347), tjIs280[pafo-1p::pafo-1::mCherry::pafo-1 3′-UTR]* (F), *control(RNAi)* (G), *leo-1(RNAi)* (H, I), *rtfo-1(RNAi)* (J), *pafo-1(RNAi)* (K), *cdc-73(RNAi)* (L), and *ctr-9(RNAi)* (M) in the hermaphrodite day 1 adult posterior gonads. (N, O) Quantitative analysis of the RNAi efficiency of *leo-1(RNAi)* (N) and *pafo-1(RNAi)* (O). *P*-values are indicated for Student’s *t*-test in comparison with the *control(RNAi)*. \*\*\**P* < 0.005. The error bars represent ± SD. (P) Percentages of oogenesis defects found in 1 day-adult wild type, mutants, transgenic rescued, *control(RNAi)*, and RNAi-knockdown animals of each PAF1C component. *P*-values are indicated for Fisher’s exact test in comparison with WT or *control(RNAi)*. \*\*\**P* < 0.005. Error bars represent ± SD. In all panels, the anterior region of the gonad was to the left, and the dorsal region was at the top of the image. The posterior gonads are shown. The orange dotted lines mark the gonad boundaries. Scale bar (white), 50 μm.

### The PAF1C is dispensable for spermatogenesis

Next, we examined whether the PAF1C is involved in spermatogenesis. When nuclei were visualized with mCherry::H2B(histone), sperm-like small cells were detected in wild type and *control(RNAi)* animals. Similarly, sperm-like cells were formed in *pafo-1(tm13447), leo-1(RNAi), pafo-1(RNAi)*, and *cdc-73(RNAi)* animals (Fig 2). Thus, these results suggest that the PAF1C is not essential for spermatogenesis.

**Fig 2.**
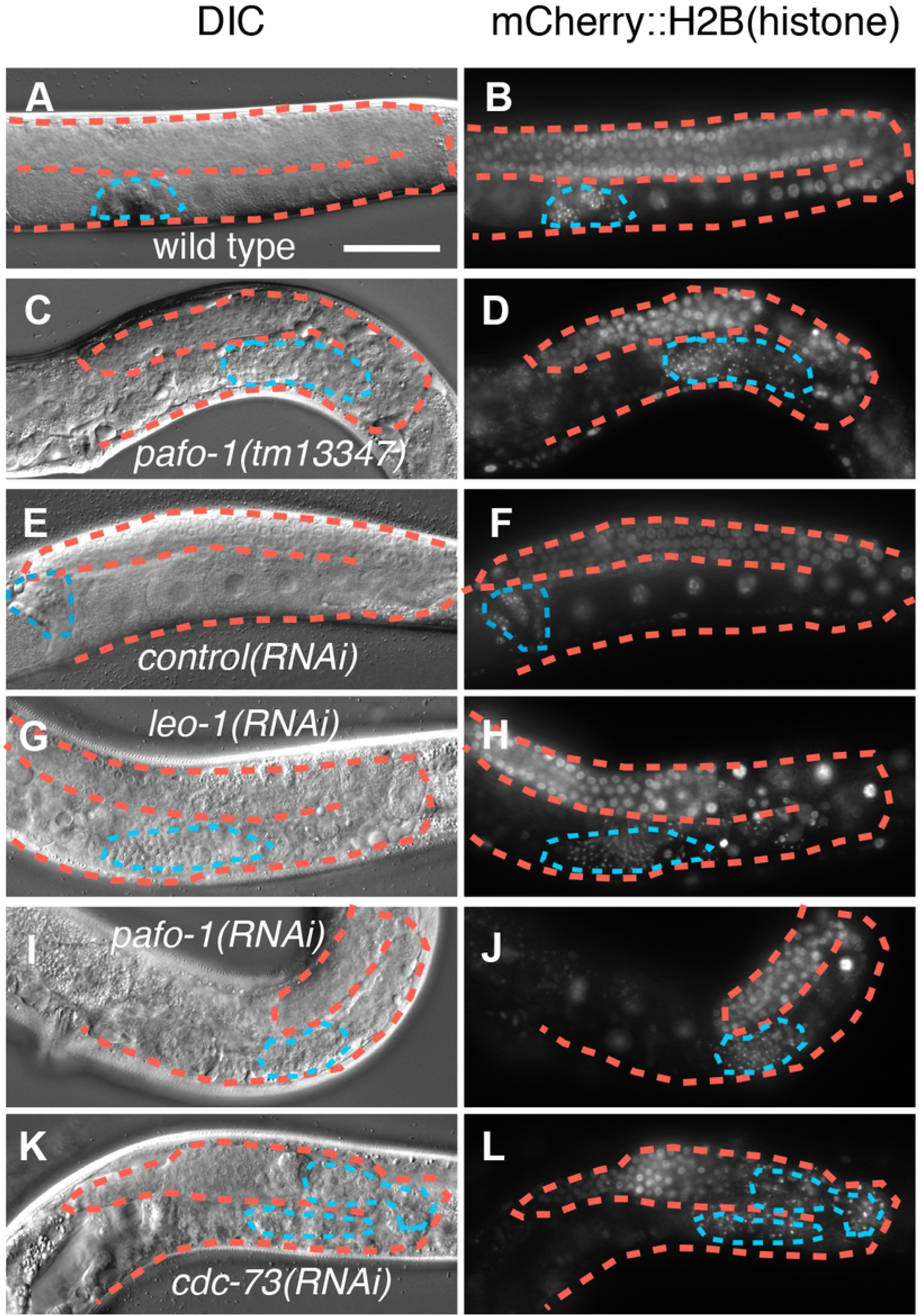
The PAF1C is Dispensable for Spermatogenesis. (A–L) Differential interference contrast (DIC) (A, C, E, G, I, and K) and fluorescence (B, D, F, H, J, and L) images of wild type (A, B), *pafo-1(tm13347)* (C, D), *control(RNAi)* (E, F), *leo-1(RNAi)* (G, H), *pafo-1(RNAi)* (I, J), and *cdc-73(RNAi)* (K, L) day 1 adult animals with *tjIs57[pie-1p::mCherry::H2B::pie-1 3′-UTR]*. In all the panels, the anterior region of the gonad is to the left, and the dorsal region is at the top of the image. The posterior gonads are shown. The orange dotted lines mark the gonad boundaries and the blue dotted lines surround the sperm-like small cells. Scale bar (white), 50 μm.

### The PAF1C is involved in the expression of OMA-1 in oocytes

Next, we investigated how the PAF1C regulates oogenesis. To visualize matured oocytes, we used an oocyte maturation marker, OMA-1::GFP, which was derived from the *oma-1* regulatory region (Fig 3A). In the day 1 adult stage of *control(RNAi)* animals, 4.1 OMA-1::GFP-positive cells were arranged linearly in the ventral region of each gonad on an average (N = 15, Fig 3B, 3C, and 3J). In contrast, the average number of OMA-1::GFP-positive cells were 1.3, 0.27, and 0.13 in *leo-1(RNAi), rtfo-1(RNAi)*, and *pafo-1(RNA)* animals, respectively (N = 15, Fig 3D–3J). These results suggest that the PAF1C positively regulates the expression of OMA-1.

**Fig 3.**
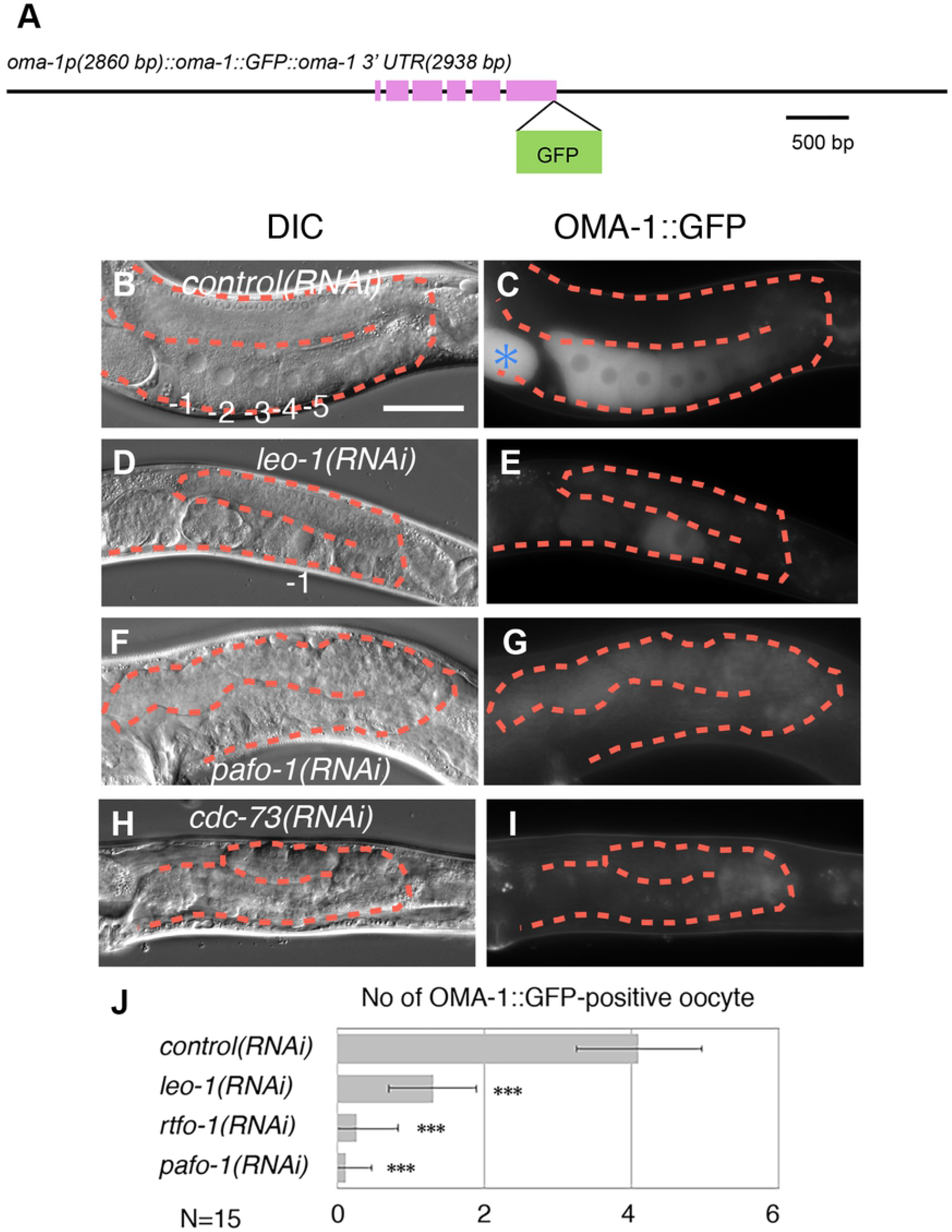
The PAF1C is Essential for the Promotion of Oocyte Maturation. (A) Genomic structure of the translational GFP-fusion construct of *oma-1*. (B–I) Differential interference contrast (DIC) (B, D, F, and H) and fluorescence (C, E, G, and I) images of *control(RNAi)* (B, C), *leo-1(RNAi)* (D, E), *pafo-1(RNAi)* (F, G), and *cdc-73(RNAi)* (H, I) day 1 adult animals with *bkcSi11[oma-1p::oma-1::GFP::oma-1 3′-UTR]* (C, E, G, and I). (J) Quantification of OMA-1::GFP-positive oocytes. *P*-values are indicated for Student’s *t*-test in comparison with *control(RNAi)*. \*\*\**P* < 0.005. The error bars represent ± SD. In all the panels, the anterior region of the gonad is to the left, and the dorsal region is at the top of the image. Posterior gonads are shown. The orange dotted lines mark the gonad boundaries. Asterisk (blue) indicates fertilized egg. Scale bar (white), 50 μm.

### Germ cell-specific expression of PAFO-1::mCherry rescues the oocyte maturation-defective phenotype of the *pafo-1(tm13447)* mutant

To determine the tissue in which the expression of PAF1C is required for oocyte maturation, we performed a tissue-specific rescue experiment using the *paflo-1(tm13447)* deletion mutant. In the day 1 adult stage of wild-type animals, approximately 4.7 OMA-1::GFP-positive cells were arranged linearly in the ventral region of each gonad on an average (N = 15, Fig 4B, 4C, and 4L). In contrast, the number of OMA-1::GFP-positive cells was significantly decreased in the *pafo-1(tm13447)* deletion mutant (the average number of OMA-1::GFP-positive cells was 0.6, N = 15; Fig 4D, 4E, and 4L). When, we introduced an integrated *tjIs280[pafo-1::mCherry]* transgene by the *pafo-1* regulatory region (Fig 4A and 4H), it almost completely rescued the oogenesis defect (the average number of OMA-1::GFP-positive cells was 4.5, N = 15, Fig 4F–H and 4L). Similarly, when we introduced integrated *pafo-1::mCherry* transgenes by germ cell-specific regulatory regions of *pie-1, bkcSi12* (Fig 4A and 4K), and *bkcSi13* (Fig 4A), they significantly rescued the oogenesis defect of the *pafo-1(tm13447)* deletion mutant (the average number of OMA-1::GFP-positive cells from *pafo-1(tm13447);bkcSi12;bkcSi11* and *pafo-1(tm13447);bkcSi13;bkcSi11* was 2.8 and 3.2, respectively, N = 15, Fig 4I–4L). These results suggest that the PAF1C regulates oogenesis in a cell-autonomous manner. Although the rescue activity with regard to the number of OMA-1::GFP-positive cells was not complete, *bkcSi13[pie-1p::pafo-1::mCherry::pie-1 3′-UTR]* rescued the sterility of the *pafo-1(tm13447)* mutant, and the *pafo-1(tm13447);bkcSi13[pie-1p::pafo-1::mCherry::pie-1 3′-UTR]* survived and produced the next generation both in the presence and absence of *bkcSi11[oma-1p::oma-1::GFP::oma-1 3’-UTR]*. Thus, germ cell expression of PAFO-1::mCherry is sufficient for the formation of functional oocytes and its maternal contribution is sufficient for embryonic, larval, larval–adult transition, and germ cell development in the next generation.

**Fig 4.**
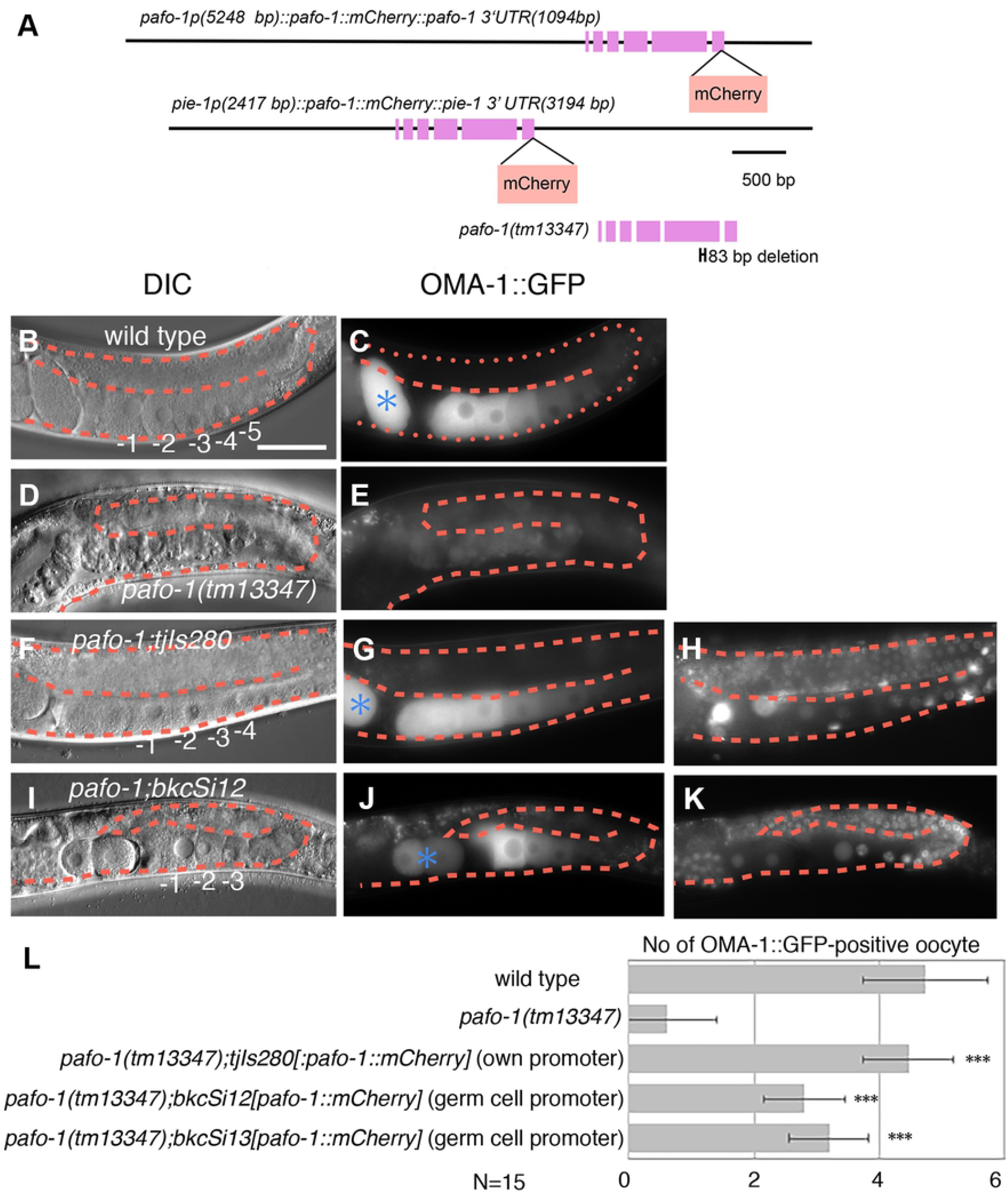
The PAF1C Promotes Oocyte Maturation in a Cell-autonomous Manner. (A) Genomic structures of the translational mCherry-fusion construct of *pafo-1*, germ cell specific mCherry-fusion construct of *pafo-1*, and the deleted region of *pafo-1(tm13447)*. (B–K) Differential interference contrast (DIC) (B, D, F, and I) and fluorescence (C, E, F, G, H, J, and K) images on wild type (B, C), *pafo-1(tm13347)* (D, E), *pafo-1(tm13347);tjIs280[pafo-1p::pafo-1::mCherry::pafo-1 3′-UTR]* (F–H), *pafo-1(tm13347);bkcSi12[pie-1p::pafo-1::mCherry::pie-1 3′-UTR]* (I–K) of day 1 adult animals with *bkcSi11[oma-1p::oma-1:GFP::oma-1 3′-UTR]*. C, E, G, and J indicate the OMA-1::GFP signals from the GFP channel. H and K indicate the *tjIs280*-derived PAFO-1::mCherry signals from the mCherry channel, and the *bkcSi12*-derived PAFO-1-mCherry signals from the mCherry channel, respectively. (L) Quantification of OMA-1::GFP-positive oocytes. *P*-values are indicated for Student’s *t*-test in comparison with *pafo-1(tm13347)*. \*\*\**P* < 0.005. The error bars represent ± SD. In all the panels, the anterior region of the gonad is to the left, and the dorsal region is at the top of the image. Posterior gonads are shown. The orange dotted lines mark the gonad boundaries. Asterisk (blue) indicates fertilized egg. Scale bar (white), 50 μm.

### Overexpression of OMA-1::GFP partially rescues the oogenesis defect in the *pafo-1(tm13447)* mutant

We tested whether the reduction in the expression of OMA-1 is the major cause of oogenesis defects in the PAF1C mutants. When OMA-1::GFP was overexpressed in the *pafo-1(tm13447)* mutant, the oogenesis defect was partially rescued (Fig 5). Therefore, a possible role of the PAF1C in the germline is to promote oogenesis by positively regulating the expression of *oma-1*.

**Fig 5.**
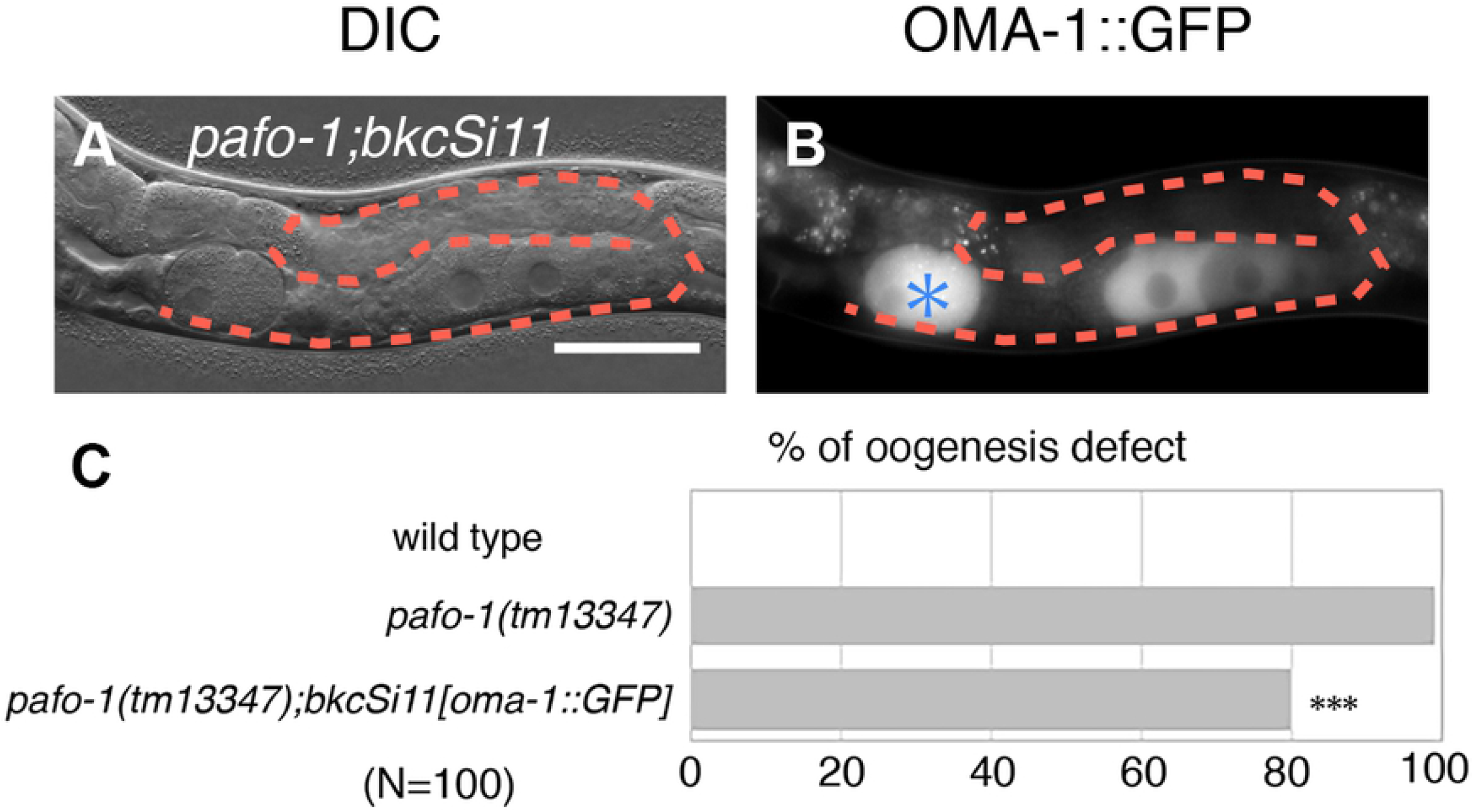
Overexpression of OMA-1::GFP Partially Rescues the Oogenesis Defect in the *pafo-1(tm13346)* Mutant. (A, B) Differential interference contrast (DIC) (A) and fluorescence (B) images of *pafo-1(tm13347);bkcSi11[oma-1p::oma-1:GFP::oma-1 3′-UTR]* day 1 adults. (C) Percentages of oogenesis defects found for 1 day-adult wild type, *pafo-1(tm13447)* mutant, and *pafo-1(tm13347);bkcSi11[oma-1p::oma-1:GFP::oma-1 3′-UTR]* animals. *P*-values are indicated for Fisher’s exact test in comparison with *pafo-1(tm13347)*. \*\*\**P* < 0.005. In all the panels, the anterior region of the gonad is to the left, and the dorsal region is at the top of the image. Posterior gonad is shown. The orange dotted lines mark the gonad boundaries. Asterisk (blue) indicates fertilized egg. Scale bar (white), 50 μm.

## Discussion

The PAF1C is a highly conserved protein complex that consists of five conserved components, LEO1, RTF1, PAF1, CDC73, and CTR9. Although it has been shown to be required in diverse biological processes, its contribution to germ cell development has not yet been explored. In this study, we performed functional analysis of the PAF1C in the germ cell development of *C. elegans* and demonstrate its requirement for cell volume expansion of oocytes and expression of OMA-1 during oogenesis.

Although the PAF1C components, LEO-1 and PAFO-1, are expressed ubiquitously, including in germ cells [20], the PAF1C is required only for oogenesis but not for spermatogenesis. Because the PAF1C is not required for sperm formation, it is unlikely that the oogenesis-defective phenotype is causative of earlier defects in germ cell development. These results suggest that PAF1C regulates a specific set of genes that are required for oogenesis.

The number of OMA-1::GFP-positive maturing oocytes was decreased in *leo-1(RNAi), rtfo-1(RNAi)*, and *pafo-1(RNAi)* animals and in *pafo-1* deletion mutants. Therefore, one possible role of the PAF1C is the promotion of OMA-1 expression. We also show that the overexpression of OMA-1::GFP partially rescued the oogenesis defect in the *pafo-1* mutant. Therefore, the PAF1C may promote oogenesis by positively regulating the specific set of downstream oocyte maturation regulators, including OMA-1 (Fig 6). Our data also suggest that the germ cell expression of PAF1C is sufficient for the formation of fully functional oocytes and that the maternal contribution of the PAF1C is sufficient for embryonic development, larval development, and larval–adult transition.

**Fig 6.**
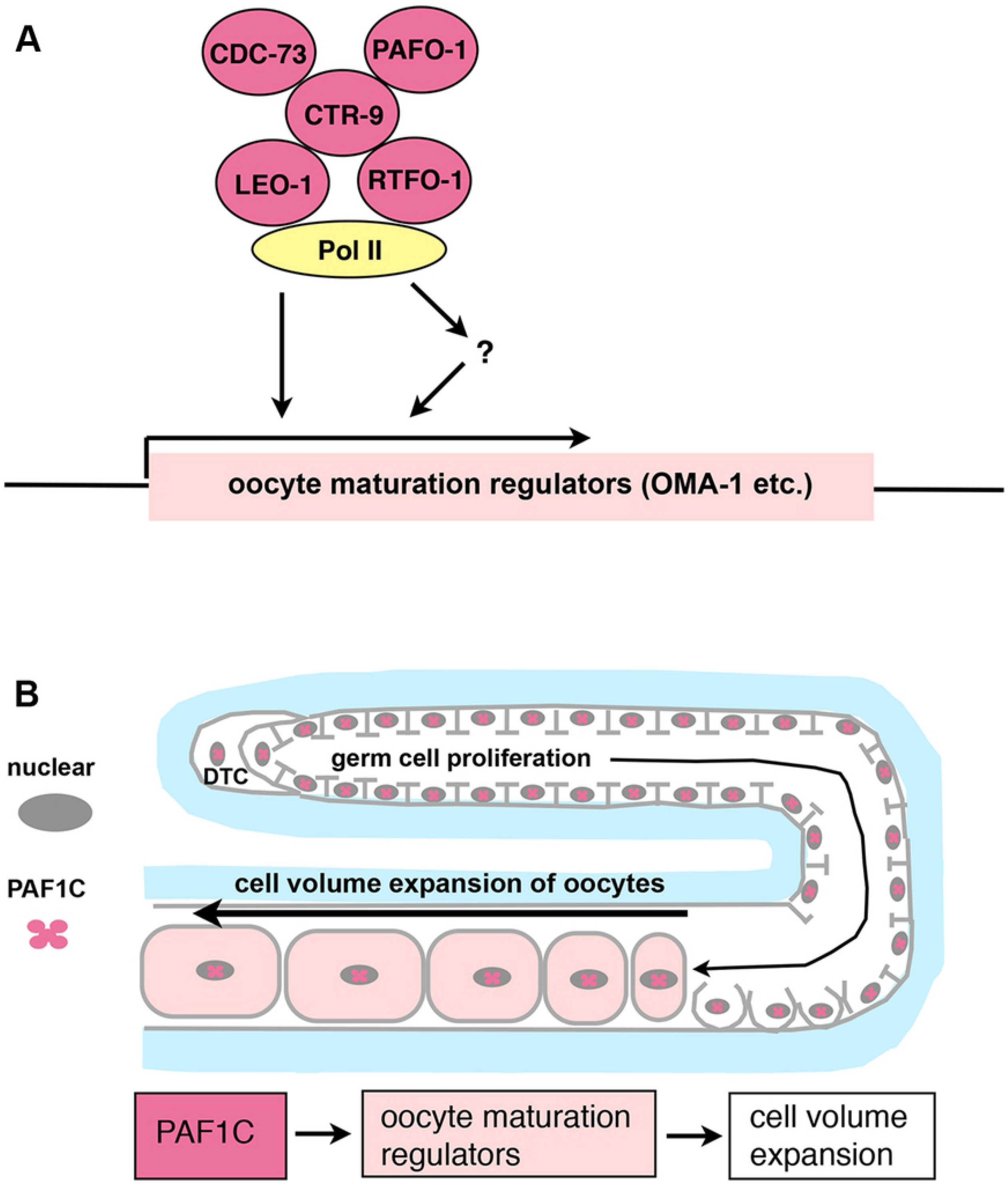
Models for the PAF1C-dependent Regulation of Cell Volume Expansion of Oocytes. (A) The PAF1C and RNA polymerase II (RNA pol II) are recruited to the regulatory region of target genes, and directly or indirectly promote the expression of the oocyte maturation regulators. (B) During oocyte maturation, the PAF1C promotes oocyte maturation regulators, which then promote cell volume expansion in the ventral region of the gonad.

It has been shown that *oma-1* and *oma-2* act redundantly to promote the later part of oocyte maturation to complete oocyte maturation [34-36]. In contrast, the cell volume expansion defect of oocytes in the PAF1C-depleted animals and mutants occurred in the early part of the oocyte maturation process. Although the overexpression of OMA-1::GFP partially rescued the cell volume expansion defect of the *pafo-1* mutant, the phenotypic similarity of the oogenesis defect was not observed in the *pafo-1 (tm13447)* mutant and the *oma-1(RNAi);oma-2(RNAi)* animals. These results indicate that the phenotypic severity of PAF1C-depleted animals was stronger than that of OMA-1/2-double depleted animals. Therefore, it is pertinent to discuss as to why the overexpression of OMA-1 rescues the cell volume expansion phenotype of the *pafo-1* mutant. A possible explanation is that although the major function of PAF1C is to promote the expression of OMA-1/2, the PAF1C may also regulate other targets that are involved in the promotion of the cell volume expansion process in parallel with the OMA-1/2 functions. Further analysis is required to confirm this hypothesis.

In this study, we found that oogenesis defects were less severe in *leo-1(RNAi)* animals among the animals with RNAi-knockdown of the five components of the PAF1C. Similar to our observations, RTF1, PAF1, CDC73, CTR9, but not LEO1, were reported to be required for the specification of an appropriate number of cardiomyocytes and for elongation of the heart tube in zebrafish [10]. Taken together, these observations suggest that in the specific context of the differentiation process, among the five PAF1C components, the requirement for LEO-1 is less critical, and this difference is conserved in vertebrates and invertebrates. At of date, there are several possible avenues for exploring this phenomenon. In yeast, Ctr9, Cdc73, and Rtf1, but not Leo1, were shown to require PAF1 at normal levels, and loss of Cdc73 resulted in a lower abundance of RTf1 [7, 37]. Therefore, it is expected that the other four components may achieve only a part of the PAF1C function in the absence of LEO-1. Although all the PAF1C components are required for its function, each component has a specific role in regulating the expression of gene encoding cell differentiation regulators, possibly by affecting the formation of the protein complex, specific protein–protein interactions, and protein– DNA/RNA interactions. Further studies are required to determine how PAF1C regulates tissue-specific development in multicellular organisms.

## Conclusion

In summary, we propose that the PAF1C promotes oogenesis in a cell-autonomous manner by positively regulating the oocyte maturation regulators, including OMA-1.

## Acknowledgements

We would like to thank Dr. Eisuke Sumiyoshi for his help with the initial phase of observation of the phenotype. We also thank Mr. Arashi Ezaki and Mr. Tasuku Hamazaki for their helpful comments. Some strains were provided by the CGC, which is funded by the NIH Office of Research Infrastructure Programs (grant number P40 OD010440), the *C. elegans* Gene Knockout Consortium, and the National Bioresource Project in Japan (led by S. Mitani).

## Supporting information

**S1 Fig.**
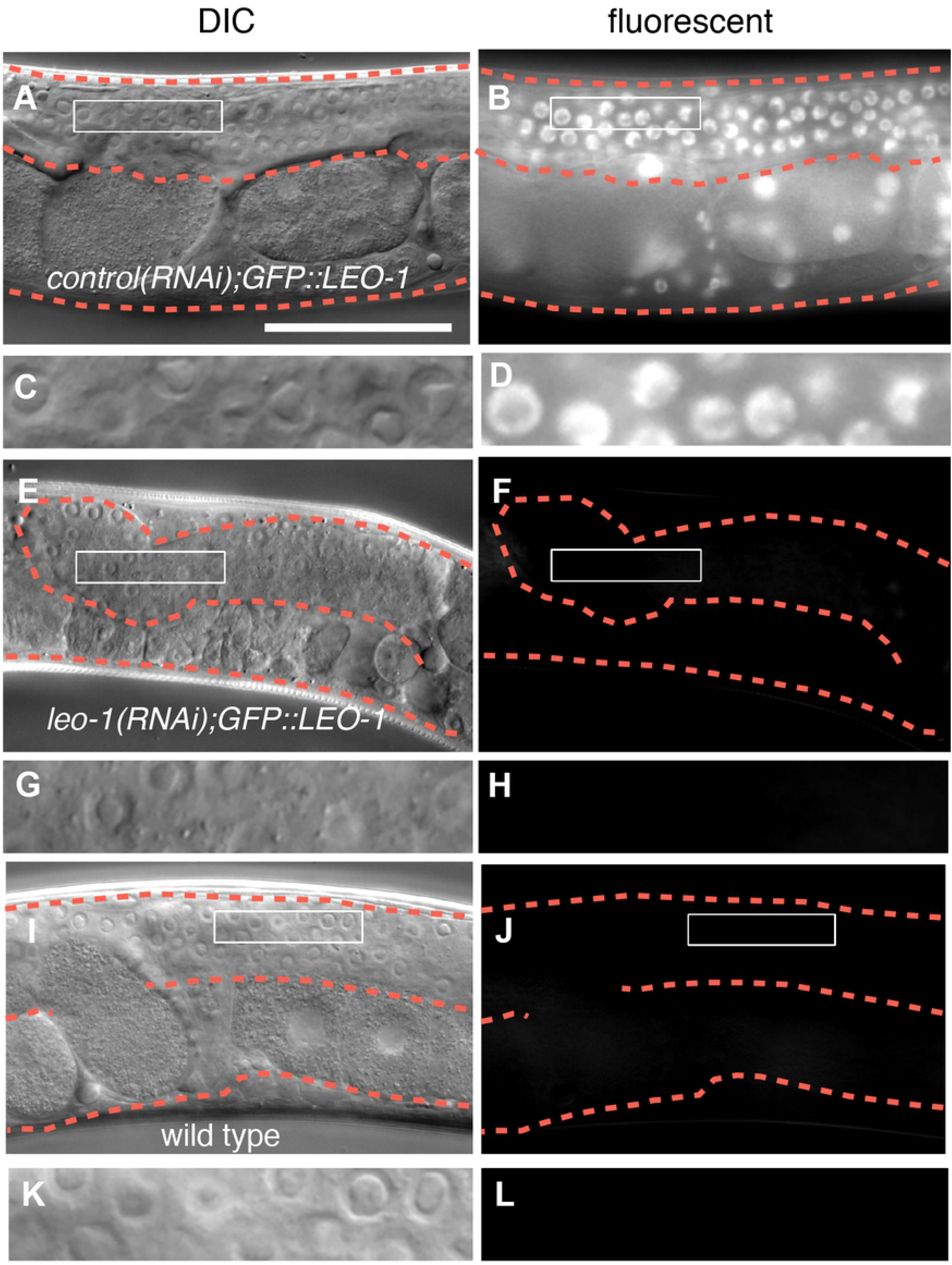
Analysis of the Efficiency of RNAi Knockdown of *leo-1*.

**S2 Fig.**
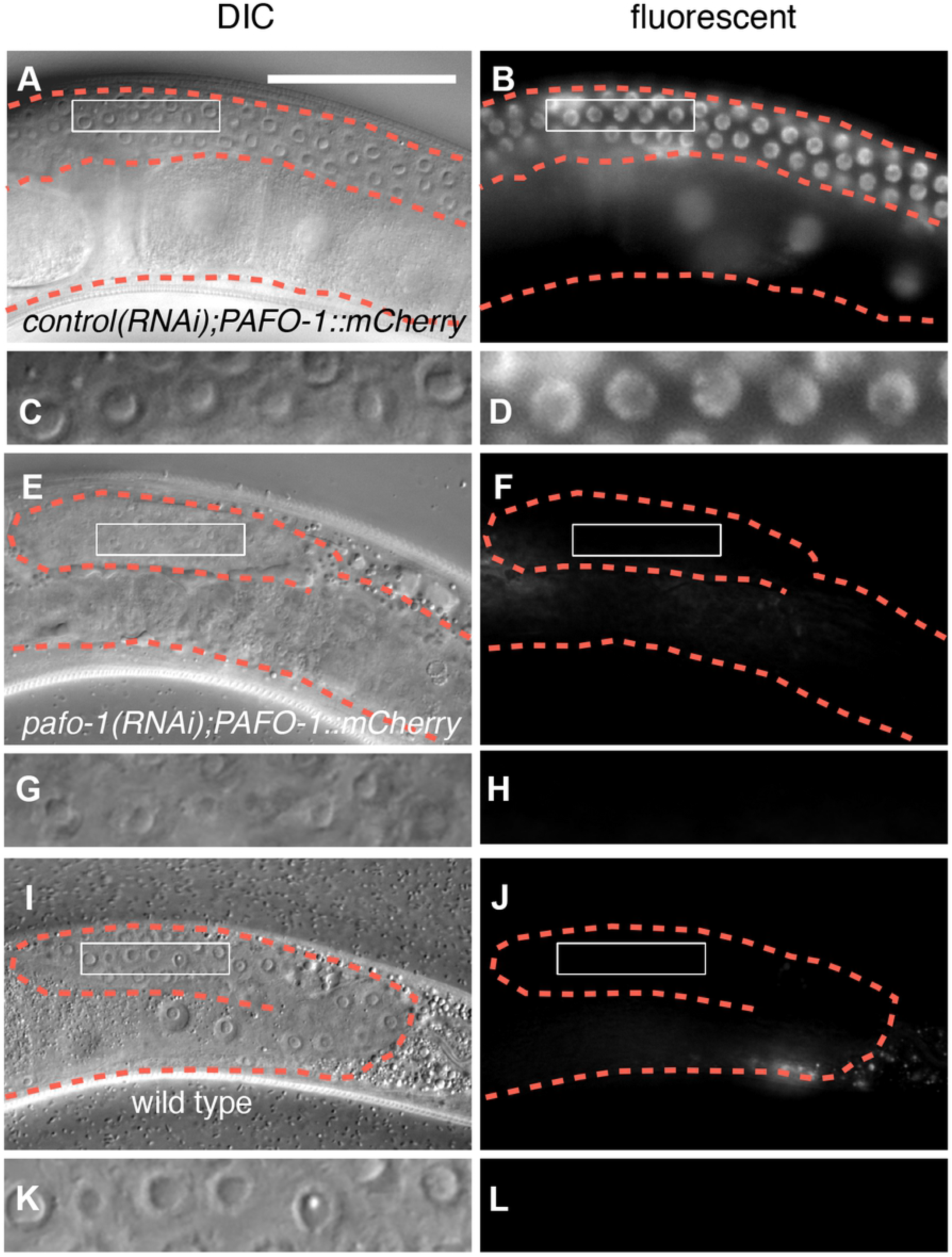
Analysis of the Efficiency of RNAi Knockdown of *pafo-1*.

**S1 Table. *Caenorhabditis elegans* Strains Constructed for this Study**.

**S2 Table. Plasmids Constructed for this Study**.

